# More fruits, more frugivores, but not necessarily more visits: findings from two tropical plant–avian frugivore communities

**DOI:** 10.64898/2026.10.08.757600

**Authors:** Rintu Mandal, Himanshu Lad, Natasha Desai, Rasika Kadam, Anand M. Osuri, Rohit Naniwadekar

**Affiliations:** Manipal Academy of Higher Education, Manipal, India; Nature Conservation Foundation, Mysuru, India; Bhavan’s College, Mumbai, India

**Keywords:** Bottom-up regulation, Carrying capacity, Frugivory, Resource dilution effect, Neutral theory, Tropical forests

## Abstract

Fruit resource availability can influence the local abundance of avian frugivores through bottom-up regulation, but whether greater resource availability translates into higher visitation rates to fruiting trees remains poorly understood. Greater fruit availability could increase visitation by supporting larger frugivore populations, or reduce visitation to individual trees if increases in frugivore abundance are not proportionate to those in fruit availability or if frugivores distribute their foraging effort. Understanding these relationships can provide insights into the relative role of abundance and trait-based mechanisms in structuring plant–frugivore communities. We compared fruit availability, avian frugivore abundance, and frugivore visitation rates on fruiting trees across two protected wet tropical forest sites: Anamalai Tiger Reserve in the Western Ghats and Namdapha Tiger Reserve in the Eastern Himalaya. Specifically, we quantified the availability of bird-consumed fruits across one-ha patches, estimated local abundances of major frugivore groups, and tested whether visitation rates on individual fruiting trees were related to frugivore abundance, body mass, and degree of frugivory. We found that small-seeded fruit availability was significantly higher in Anamalai than in Namdapha. Local abundances of several relatively small-bodied frugivore groups, including bulbuls, barbets, fairy-bluebirds, and mynas, were also higher in Anamalai. However, visitation rates of several frugivore groups on individual fruiting trees were higher in Namdapha. Visitation rates across all avian frugivore species were associated with frugivore abundances only in Anamalai, and not associated with body mass or degree of frugivory in either site. Thus, although greater fruit availability was likely associated with higher frugivore abundance, this increase in abundance did not consistently translate into higher visitation rates at individual fruiting trees, which provides some support for the resource dilution hypothesis. Our results highlight the importance of considering fruit resource availability and frugivore abundance when examining variation in tropical plant-frugivore interactions.

## INTRODUCTION

Understanding processes that influence interactions between species is central to community ecology, as interaction strengths may underpin many ecosystem functions (Duffy, 2002; Schleuning et al., 2015). In tropical forests, interactions between fruiting plants and vertebrate frugivores determine seed dispersal of fleshy-fruited species, influencing their patterns of recruitment, spatial structure, and ultimately the maintenance of tree diversity (Bascompte & Jordano, 2007; Sengupta, 2022). Considerable attention has been devoted to understanding the role of plant attributes such as morphological and chemical properties of fruits, fruit crop size, and fruiting neighbourhoods, and frugivore attributes, such as gape size, in influencing these interactions (Blendinger et al., 2008; Burns, 2013; Guerra et al., 2017; Chenevert et al., 2026). By contrast, comparatively less is known about the influence of landscape-scale variation in resource availability on frugivore abundance and plant–frugivore interaction frequencies.

Resource availability can influence consumer abundance through bottom-up regulation of ecosystem carrying capacity (Tilman, 1982; Polis, 1999). In the context of the frugivory literature, this has been examined through the lens of the fruit resource tracking hypothesis, where patches with higher fruit availability have been shown to harbour higher abundances of frugivores (Rey, 1995; Burns, 2004; Blendinger et al., 2012). These studies suggest that fruit availability is a major driver of local frugivore abundance. Now, if interaction frequencies scale primarily with consumer abundance, higher resource availability should support more frugivores in the local landscape and consequently, greater visitation rates on fruiting trees. Alternatively, in accordance with the resource dilution hypothesis, greater resource availability may reduce visitation rates on individual fruit trees despite supporting higher frugivore abundances, as a likely outcome of consumers distributing their foraging efforts across a large pool of available resources (Otway et al., 2005). Such decoupling between frugivore abundance and visitation rates has often been observed at finer spatial scales of fruiting neighbourhoods, where high neighbourhood fruit availability can reduce visitation rates or fruit removal from individual trees (Saracco et al., 2005; Smith & McWilliams, 2014; Hopson et al., 2020; Chenevert et al., 2026). Conversely, when fruit availability is low, frugivores may concentrate their foraging activity on the limited fruiting trees available, potentially increasing visitation rates on individual fruiting trees (Howe, 1977; Ragusa-Netto, 2002). Whether similar dilution processes operate at a broader landscape or biogeographic scales remains largely unknown. At small spatial scales, frugivores could easily redistribute their activity among individual trees, whereas at broader scales, differences in resource availability may influence frugivore densities and movements. Therefore, greater resource availability at larger scales could result in higher frugivore densities and potentially provide a larger quantum of fruit resources, thereby resulting in different relationships between resource availability, frugivore abundance, and visitation patterns than those observed at local fruiting neighbourhoods.

In addition to resource availability and frugivore abundance, frugivore species attributes such as body size and degree of frugivory may also influence their visitation rates on fruiting trees, yet their influence remains relatively poorly understood (but see Peña et al., 2023). As per neutral theory, visitations of frugivores with fruiting plants should largely reflect their encounter probabilities and therefore species abundance (Donoso et al., 2017; Pizo et al., 2022). By contrast, visitation rates may be influenced by niche-based processes related to frugivore traits. For example, large-bodied frugivores are typically rarer in communities than small-bodied frugivores (White et al., 2007; Velho et al., 2012), potentially resulting in relatively fewer fruiting tree visits by the former. Similarly, obligate frugivores can be expected to show higher visitation frequencies on fruiting trees, as a consequence of their greater dependence on fruits, and they might track fruit resources more closely than facultative frugivores (Pizo et al., 2021; Morán-López et al., 2025). Thus, examining whether species-level visitation is explained by abundance alone, or additionally by attributes related to body size and dietary dependence, can help identify factors influencing plant-frugivore interactions.

We conducted a comparative study across two sites in South Asia: Anamalai Tiger Reserve in the Western Ghats, and Namdapha Tiger Reserve in the Eastern Himalaya. While the two sites harbour similar wet tropical forests (Proctor et al., 1998; Muthuramkumar et al., 2006), they differ in the diversity of their avian frugivore species pools and resource availability. This enables us to examine how resource availability, avian frugivore abundance, and visitation patterns are linked across differing ecological contexts.

To this end, we compared the availability of bird-consumed ripe fruits across fruiting patches between the two sites. We categorised plant species into small-, medium-, and large-seeded plant species, as seed size can significantly influence the diversity and composition of visiting frugivores (Naniwadekar et al., 2025). Next, we examined how the local abundance of major avian frugivore groups and their visitation rates to individual fruiting trees varied between the two sites. Finally, we tested how frugivore abundance and two key frugivore functional traits: body mass and degree of frugivory, influenced their visitation frequencies to individual fruiting trees. Specifically, our predictions were: 1) Frugivore abundance would be higher in the site with greater fruit availability, following expectations of carrying capacity through bottom-up regulation. 2) We considered two alternative predictions for visitation rates. If increases in frugivore abundance are proportionally greater than increases in fruit availability, visitation rates per fruiting tree should be higher at the site with greater frugivore abundance. Conversely, if fruit availability increases disproportionately relative to frugivore abundance, frugivores may distribute their foraging effort across a larger resource pool, resulting in lower visitation rates per fruiting tree, as predicted by the resource dilution hypothesis. 3) Visitation rates of individual frugivore species are jointly influenced by their local abundance and key functional traits (body mass and degree of frugivory). By comparing fruit resource availability, frugivore abundance, and visitation rates across contrasting tropical forests, we test whether differences in resource availability and frugivore abundance influence frugivore visitation rates at fruiting trees, and whether variation in visitation among frugivore species is associated with body size and degree of frugivory. Our study aims to identify mechanisms linking consumer populations to the strength of ecological interactions.

## MATERIALS AND METHODS

### Study area

We conducted the study in two tropical forest sites in India: Anamalai Tiger Reserve (10°12′–10°35′N, 76°49′–77°24′E; 650–1,000 m a.s.l.) in the Western Ghats and Namdapha Tiger Reserve (27°23′–27°39′N, 96°15′–96°58′E; 550–800 m a.s.l.) in the Eastern Himalaya. Both sites fall within the “Tropical and Subtropical Moist Broadleaf Forests” biome (Olson et al., 2001). The study was conducted within an approximately 15 km^2^ contiguous patch of evergreen forests within the larger Protected Area. Sampling locations represented lowland wet evergreen forests within protected areas and lacked any recent history of anthropogenic disturbance. Both sites receive approximately 2,500–3,000 mm of rainfall annually and harbour lowland wet tropical evergreen forests (sensu Whitmore, 1984). By sampling across two sites using identical methods, we could get data from independent locations with differing diversity of avian frugivores.

In Anamalai, long-term phenology monitoring indicates that fruiting of bird-consumed tree species peaks between April and June (Madhavan et al., 2026). Comparable long-term phenological information is unavailable for Namdapha. However, fruiting of bird-consumed species in Pakke Tiger Reserve, another lowland wet evergreen forest site in the Eastern Himalaya, peaks between March and June (Datta et al., 2025), suggesting broadly similar phenological patterns in the region. These sites share several major lineages of fleshy-fruited plants, including Myristicaceae, Lauraceae, Meliaceae, Moraceae, and Annonaceae, and avian frugivore groups such as Bucerotidae, *Psilopogon* spp., Pycnonotidae, *Gracula* spp., *Ducula* spp., and *Treron* spp. However, Anamalai (26 avian frugivore species) is relatively species-poor compared to Namdapha (40 frugivore species), as the Western Ghats have lower within-clade diversity than the Eastern Himalaya (see Naniwadekar et al., 2025; Mandal et al., in press). This provides a useful basis for comparing plant-frugivore assemblages across two geographically distinct tropical forest assemblages with similar environments and nested frugivore assemblages (Naniwadekar et al., 2025).

### Fruit availability at the local scale

To estimate the availability of bird-consumed ripe fruits, we monitored 14 fixed-length transects (each 500 m long) in Namdapha between November 2022 and April 2023 and 13 transects in Anamalai from November 2023 to April 2024. Transects were laid by minimally clearing overhanging vegetation to allow passage and were georeferenced using a Garmin GPSMAP 78S. We monitored each transect throughout the study period, with seven sampling occasions in Namdapha and 10 in Anamalai, resulting in a total sampling effort of 49 km in Namdapha and 65 km in Anamalai.

We estimated the abundance of ripe fruits on bird-consumed woody plants within 10 m on either side of each transect, resulting in a 1-ha sampling area per transect (500 × 20 m^2^). Fruit abundance on individual fruiting woody plants was estimated following the approaches outlined in previous studies (Davidar & Morton, 1986; Naniwadekar et al., 2019). The method was adjusted according to fruit size, fruit crop size, and canopy structure. For trees bearing relatively small crops of large fruits (e.g., *Myristica* spp.), all ripe fruits were counted directly. For trees bearing larger crops, ripe fruits were counted in representative portions of the canopy and extrapolated to the entire crown. Depending on canopy structure, representative portions comprised a defined fraction of the crown (e.g., one-quarter of the canopy) or multiple branches distributed across the crown. For species bearing very large crops, such as *Ficus* spp., ripe fruits were counted on multiple representative branches and extrapolated to the entire canopy based on the distribution of fruits. A plant species was considered bird-consumed if at least one frugivory event, defined as consumption of fruit by an avian frugivore, was observed on that species during the study period. Only fruits from plant species meeting this criterion were included in estimates of bird-consumed fruit availability and used for further analysis.

### Local abundance of avian frugivores

We estimated encounter rates of avian frugivores by monitoring the same transects used to estimate fruit availability and during the same sampling sessions described above. Transects were walked at a slow pace (mean: 50 min per 500 m; range: 29–84 min), and frugivore detections were recorded visually and acoustically. For visual detections, we recorded the frugivore species identity and number of individuals. For auditory detections, we recorded species identity. Detections beyond 100 m on either side of the transect were excluded from analyses. Transect walks were conducted on clear days and were avoided during rainfall to minimise weather-related differences in detectability. In Namdapha, transects were surveyed both in the morning and evening, whereas in Anamalai they were sampled only in the morning because of increased risk of elephant encounters under low-light conditions. We started morning transect walks approximately one hour after sunrise at both sites, while evening surveys in Namdapha began approximately one hour before sunset, corresponding to periods of relatively high frugivore activity reported from the region (Naniwadekar et al., 2019). Additionally, we did not find differences in mean encounter rates of frugivores between transects walked in mornings and evenings in Namdapha (Fig. S1). A bird species was classified as a frugivore at a given study site if at least one individual was observed consuming fruit during the study period.

### Frugivore visitations on fruiting trees

We conducted focal tree watches on fruiting trees from November 2022 to April 2023 in Namdapha and from November 2023 to May 2024 in Anamalai. We searched for bird-consumed fruit tree species bearing ripe fruits along existing paths and trails within the study areas. At each focal tree, two observers conducted a six-hour focal watch using 8 × 40 or 8 × 42 binoculars, following established protocols (Naniwadekar et al., 2025). During each watch, we recorded the identity and number of avian frugivores arriving at the focal tree. Birds were considered frugivores on a given focal tree species only when they consumed at least one fruit of that particular species.

### Fruit morphology and frugivore functional traits

We measured at least five ripe fruits from each bird-consumed fruiting tree species recorded during transect surveys or focal watches at each study site. To compare ripe fruit availability for each size category between our study sites, we categorised plant species into three seed-size categories based on seed width: small-seeded (seed width < 5mm), medium-seeded (5–15 mm) and large-seeded (> 15 mm), following Naniwadekar et al. (2019). Body mass (g) and dietary information (percentage of fruit in the diet) for avian frugivore species were obtained from the global bird trait databases ‘Avonet’ and ‘EltonTraits’, respectively (Wilman et al., 2014; Tobias et al., 2022).

### Analyses

#### Abundances of bird-consumed fleshy fruits

For each 1 ha (500 × 20 m^2^) transect patch and sampling occasion, we summed the number of ripe fruits across all bird-consumed plant species separately for small-, medium-, and large-seeded species. Because fruit abundance varied by several orders of magnitude among plant species and patches, we natural log-transformed values prior to analysis. We compared fruit abundance between the two study sites using a linear mixed-effects model, with log-transformed fruit abundance as the response variable and study site as a fixed effect, using the R package ‘lme4’ (Bates et al., 2015). We carried out separate analyses for each seed size category, as comparing fruit availability across seed size classes was not of interest to us, as small-seeded fruits are known to be more abundant than large-seeded ones (Jordano, 2000; Dombroskie et al., 2016). To account for temporal replication, we incorporated transect-patch identity as a random intercept. Pairwise contrasts were examined using the R package ‘emmeans’ (Lenth et al., 2026). For graphical presentation, we calculated mean and bootstrapped 95% confidence intervals (with 1,000 iterations) of log-transformed fruit abundance across transect patches within each site for different seed size categories.

### Avian frugivore abundance

We compared the abundance of six major frugivore groups: Pycnonotidae (bulbuls), Irenidae (fairy-bluebirds), Megalaimidae (barbets), Sturnidae (mynas and starlings), Columbidae (pigeons), and Bucerotidae (hornbills) across the two sites. These groups are well represented across both sites. Past studies have demonstrated the critical role of these groups in frugivory and seed dispersal in Asian tropics (Naniwadekar et al., 2019; Gopal et al., 2020). These groups comprised 96% of visits on fruiting trees in Anamalai and 64% in Namdapha. Local abundance was quantified as encounter rate, defined as the number of flocks detected per km of transect surveyed. We preferred estimating flock encounter rates, since 75% of our detections on trails at each site were call-based, where it was not possible to determine the number of individuals. Flock encounter rates were strongly correlated with individual encounter rates for the selected frugivore groups (Pearson’s *r* = 0.87, *p* < 0.001; Fig. S2). To test whether flock encounter rates differed between study sites within each frugivore group, we fitted separate negative-binomial generalised linear mixed-effects models (GLMMs) for each group, with study site as a fixed effect and transect identity as a random intercept to account for temporal replications. For graphical presentation, we calculated the mean encounter rate and bootstrapped 95% confidence intervals (with 1,000 iterations) across transects within each site for each frugivore group.

### Avian frugivore visitations

We compared visitation rates between sites for the same six avian frugivore groups. For each group, we calculated bird visitation rate on each focal tree species at each site by dividing the total number of individuals recorded during focal watches of that tree species by the corresponding total observation effort (hours). Because visitation rates were strongly right-skewed, they were log-transformed before analysis to improve the normality and homoscedasticity of model residuals. To test whether visitation rates differed between sites, we fitted a separate linear model for each frugivore group, with log-transformed visitation rate as the response variable and site as the predictor. For graphical representation, we also calculated the mean visitation rate and bootstrapped 95% confidence intervals (with 1000 iterations) across focal tree species within each site for each frugivore group.

### Frugivore visitations and their local abundance and functional traits

Using all avian frugivore species recorded at a site, we examined whether interspecific variation in visitation rates was associated with their local abundance and functional traits. We fitted a separate linear model for each study site, with frugivore species as the observational units. Specifically, we modelled the overall mean visitation rate (number of birds hour^−1^) of each frugivore species across fruiting species at a site as a function of its mean encounter rate on transects (number of detections km^−1^), body mass (g), and frugivory degree (percentage fruit in the diet). Visitation rate was log-transformed to improve the distributional properties of the model residuals. All predictors were standardised to a mean of zero and a standard deviation of one before model fitting. We assessed multicollinearity among predictors using variance inflation factors calculated with the R package ‘car’ (Fox et al., 2026), and evaluated model explanatory power using adjusted *R^2^*.

## RESULTS

We recorded ripe fruits of 18 bird-consumed fleshy-fruited plant species at each site (Anamalai and Namdapha), based on frugivory interactions recorded either during focal watches or opportunistically. Transect surveys recorded 25 avian frugivore species in Anamalai and 35 in Namdapha (Table S1). During focal tree watches, we recorded frugivory interactions involving 26 bird species in Anamalai and 40 in Namdapha, across 28 fruiting tree species at each site. For comparisons of frugivore abundance and visitation rates, we focused on six major frugivore groups (Pycnonotidae, Irenidae, Megalaimidae, Sturnidae, Columbidae, and Bucerotidae). These groups were common to both sites.

### Abundances of bird-consumed fleshy fruits

The linear mixed-effects models suggested that mean ripe fruit abundance differed between the two study sites for small-seeded plant species, but not for medium- or large-seeded species (Fig. 1; Table S2,3). For small-seeded species, mean (95% CI) log-transformed ripe fruit abundance in one-ha patches was higher in Anamalai (3.91 (2.97–4.94) fruits ha^−1^) than in Namdapha (2.15 (1.38–3.18)). On an absolute scale, these estimates corresponded to approximately a six-fold difference in mean fruit abundance between sites. Mean (95% CI) ripe fruit abundance of medium-seeded bird-consumed plants (Anamalai: 1.90 (0.84–3.21) fruits ha^−1^, Namdapha: 0.97 (0.72–1.22) fruits ha^−1^) and large-seeded species (Anamalai (1.26 (0.77–1.70) fruits ha^−1^) and Namdapha (0.96 (0.57–1.40) fruits ha^−1^) did not differ significantly between the two sites. (Fig. 1; Table S2,3).

**Figure 1.**
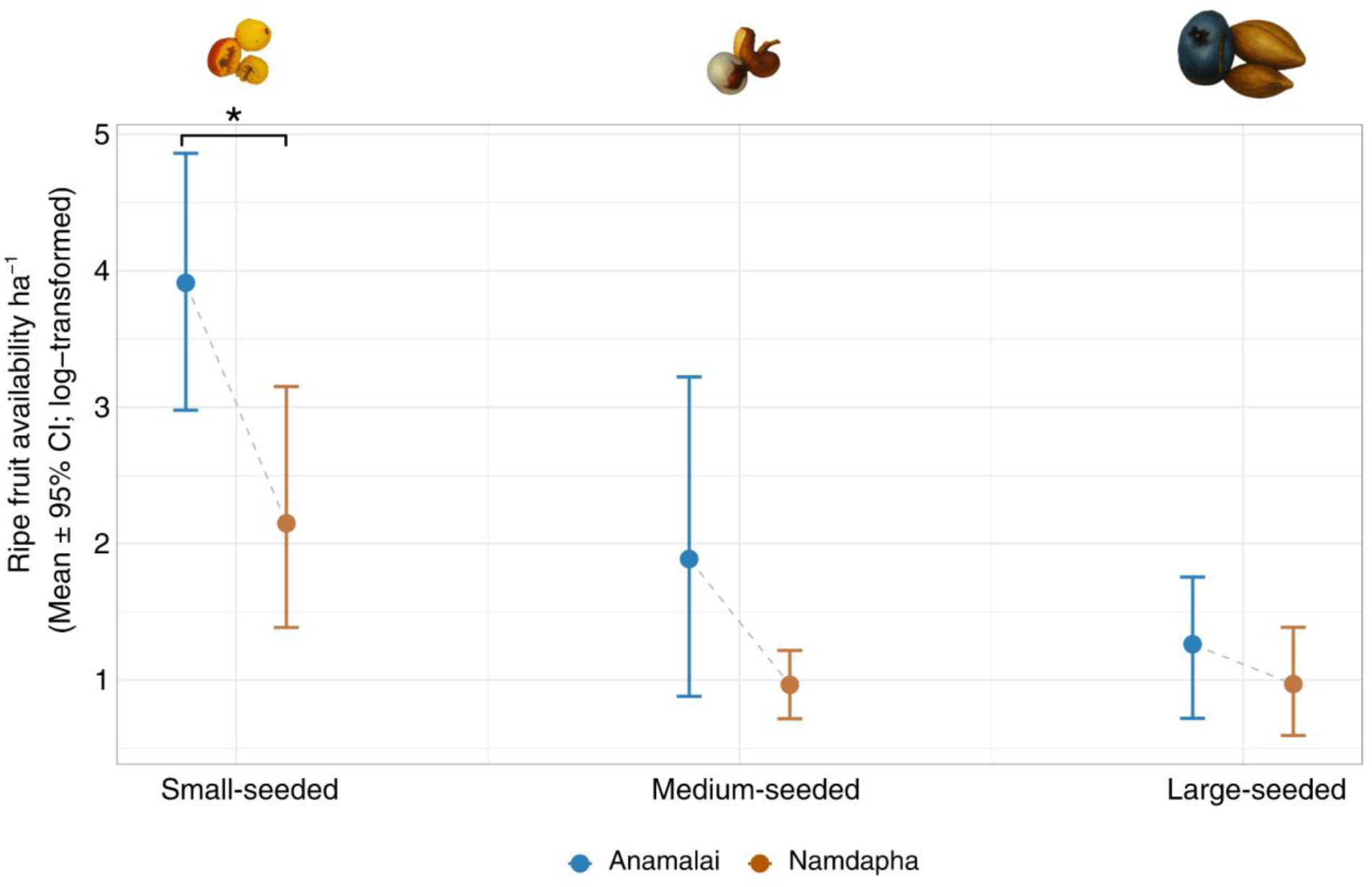
Mean (bootstrapped 95% confidence interval) natural log-transformed ripe fruit availability per hectare for small- (e.g. *Ficus*), medium- (e.g. *Trichilia*), and large-seeded (e.g. *Canarium*) bird-consumed fleshy fruited plants in Anamalai and Namdapha. Asterisk indicates significant differences between sites (*p* < 0.05).

### Avian frugivore abundance

Frugivore encounter rates (detections km⁻¹) were two- to seven-times higher in Anamalai than in Namdapha for four of the six major frugivore groups examined (Fig. 2A). Negative-binomial generalised linear mixed-effects models followed by post-hoc pairwise comparisons indicated significantly higher mean encounter rates in Anamalai than in Namdapha for the relatively smaller-bodied Pycnonotidae (5.15 (bootstrapped 95% CI: 4.51–5.80) vs. 2.23 (1.86–2.57)), Irenidae (2.57 (2.17–2.95) vs. 0.78 (0.47–1.12)), Megalaimidae (7.75 (6.62–9.06) vs. 2.96 (2.53–3.43)), and Sturnidae (2.43 (1.83–2.94) vs. 0.33 (0.16–0.53)) (Table S4). In contrast, encounter rates did not differ significantly between sites for the relatively larger-bodied Columbidae (0.40 (0.22–0.59) in Anamalai vs. 0.53 (0.35–0.74) in Namdapha) or Bucerotidae (1.89 (1.57–2.28) vs. 2.02 (1.57–2.45)) (Table S4).

**Figure 2.**
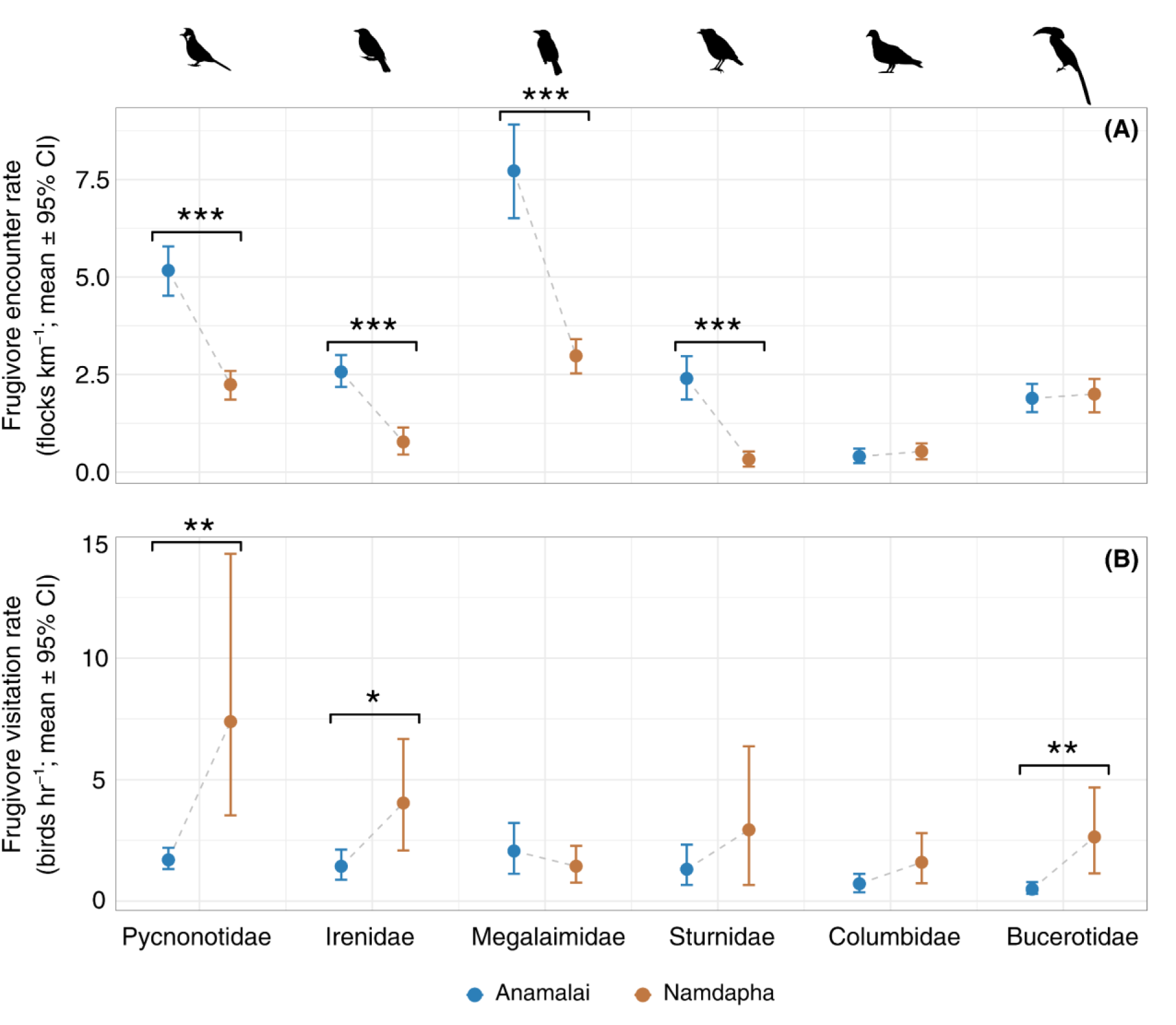
Mean (bootstrapped 95% confidence intervals) (A) encounter rates (detections km^-1^) and (B) visitation rates (number of birds hr^-1^) for six major avian frugivore groups common to Anamalai and Namdapha. Asterisks indicate significant pairwise differences between sites (*p < 0.05*, ** *p* < 0.0, and \*\*\**p* < 0.001). Bird silhouettes representing the respective frugivore groups were obtained from http://www.phylopic.org.

### Avian frugivore visitations on fruiting trees

Visitation rates (number of birds hour^−1^) showed a contrasting pattern for several frugivore groups (Fig. 2B). Linear model results showed that mean visitation rates were significantly higher in Namdapha than in Anamalai for Pycnonotidae (7.40 (95% CI: 3.52–14.10) vs. 1.69 (1.30–2.18)), Irenidae (4.07 (1.92–6.80) vs. 1.43 (0.84–2.04)), and Bucerotidae (2.62 (1.14–4.50) vs. 0.50 (0.29–0.78)) (Table S5). We did not find significant differences between the two sites for Megalaimidae (1.40 (0.77–2.21) in Namdapha vs. 2.05 (1.07–3.26) in Anamalai), Sturnidae (2.95 (0.71–6.69) vs. 1.33 (0.67–2.28)) or Columbidae (1.61 (0.67–2.77) vs. 0.72 (0.38–1.15)) (Table S5).

### Frugivore visitations and their local abundance and functional traits

Linear model results showed that frugivore visitation rates on fruiting trees were positively associated with frugivore abundance in Anamalai (*p* = 0.003; Fig. 3A; Table 1). However, such a relationship didn’t hold in Namdapha (Fig. 3D; Table 1). Additionally, neither degree of frugivory nor body mass of avian frugivores was significantly associated with visitation rates in either site (Fig. 3; Table 1). The model explained a substantial amount of variation in Anamalai (adjusted *R^2^*= 0.28). Multicollinearity among predictors was low in both models (all VIFs < 1.5), indicating that encounter rate, body mass, and degree of frugivory contributed largely independent information (Table S6).

**Figure 3.**
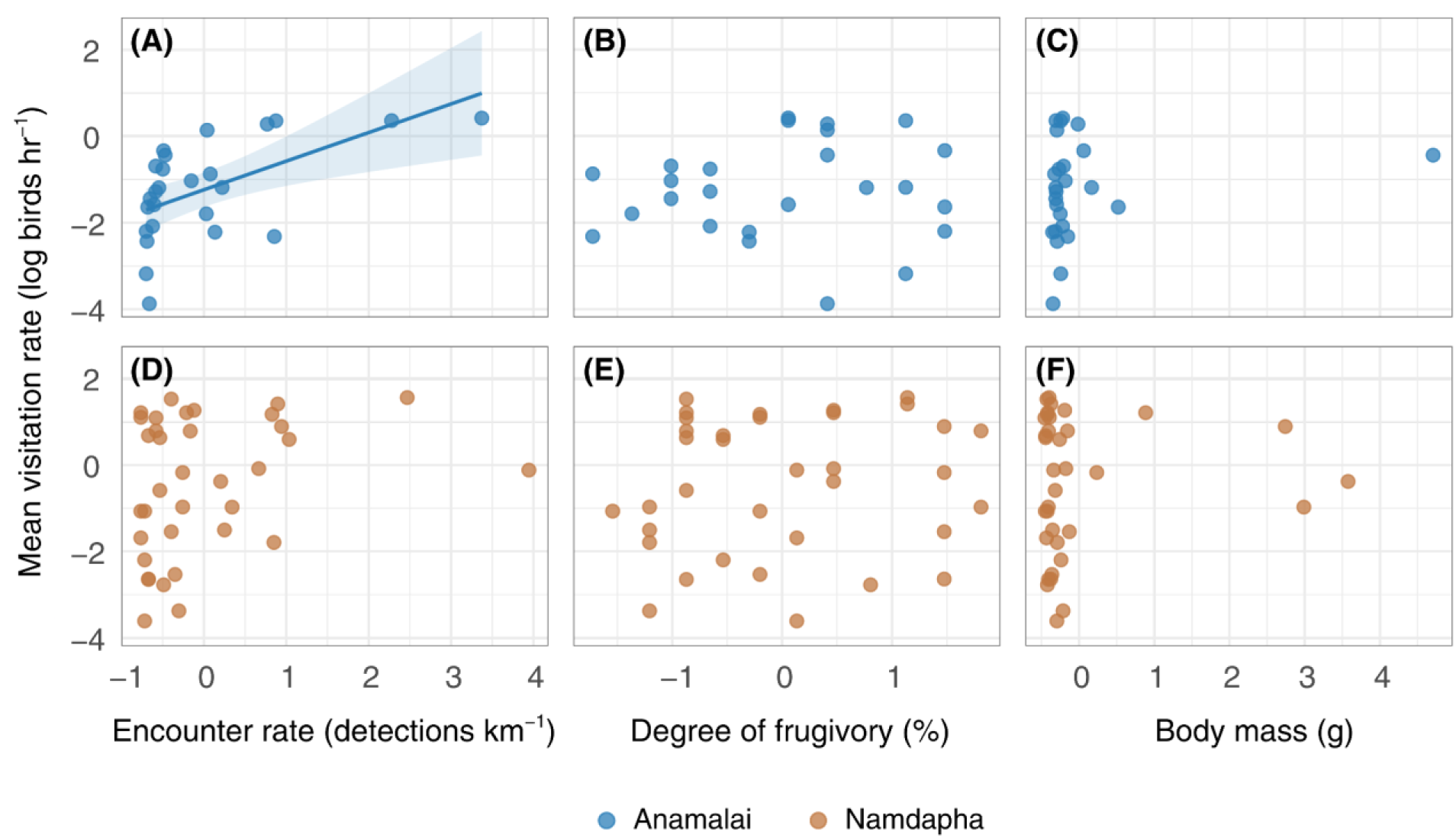
Relationships between frugivore visitation rates and frugivore attributes in Anamalai and Namdapha. Panels show log-transformed mean visitation rate (birds h⁻¹) in relation to standardised mean encounter rate (detections km⁻¹) in Anamalai (A) and Namdapha (D), standardised degree of frugivory (percentage of fruit in the diet) in Anamalai (B) and Namdapha (E), and standardised body mass (g) in Anamalai (C) and Namdapha (F). Predictors were standardised separately within each site to a mean of zero and a standard deviation of one. The solid line and shaded band in panel A represent the fitted linear-model relationship and its 95% confidence interval, respectively, with the other predictors held at their site-specific means. The fitted relationship is shown only where it was statistically significant (*p* < 0.05). Each point represents a frugivore species.

**Table 1.** Results of site-specific linear models examining associations between visitation rates and the local abundance and functional traits of all avian frugivore species recorded in Anamalai and Namdapha. The response variable was the log-transformed mean visitation rate of each frugivore species across focal fruiting-tree species. Encounter rate, degree of frugivory, and body mass were standardised separately within each site to a mean of zero and a standard deviation of one. Predictor estimates therefore represent the expected change in log-transformed visitation rate associated with a one-standard-deviation increase in the predictor, with the other predictors held constant. SE denotes standard error. Statistically significant effects (*p* < 0.05) are shown in bold.

| Site | Predictor | Estimate | SE | *p*-value |
| --- | --- | --- | --- | --- |
| Anamalai | Intercept | -1.24 | 0.19 | <0.001 |
| Anamalai | **Encounter rate** | **0.66** | **0.20** | **0.003** |
| Anamalai | Degree of frugivory | 0.09 | 0.20 | 0.675 |
| Anamalai | Body mass | 0.25 | 0.20 | 0.233 |
| Namdapha | Intercept | -0.45 | 0.27 | 0.100 |
| Namdapha | Encounter rate | 0.42 | 0.28 | 0.137 |
| Namdapha | Degree of frugivory | 0.07 | 0.31 | 0.823 |
| Namdapha | Body mass | 0.03 | 0.30 | 0.921 |

## DISCUSSION

We compared the abundances of fleshy fruits and avian frugivores, and their interaction strengths across two wet tropical forest sites that differed in avian frugivore species richness. Consistent with the expectations of bottom-up regulation, Anamalai, the site that had greater local availability of small-seeded fruits, supported higher abundances of several major frugivore groups than Namdapha. Interestingly, however, greater frugivore abundance in Anamalai did not translate into higher visitation rates on individual fruiting trees. Instead, visitation rates of several frugivore groups were higher in Namdapha. Finally, visitation rates of individual frugivore species on focal trees were associated with their local abundance only in Anamalai. On the other hand, frugivore functional traits, such as degree of frugivory and body mass, showed no significant association with their visitation rates. Together, these results suggest that although fruit availability and frugivore abundance were positively associated across sites, greater fruit availability and frugivore abundance did not consistently translate into higher visitation rates at fruiting trees.

In line with the predictions of bottom-up regulation and carrying capacity, we found frugivore abundances to be higher in Anamalai, the site with greater local fruit availability. Small-seeded fruits, which form a substantial component of the diets of many tropical avian frugivores (Wheelwright, 1985; Naniwadekar et al., 2025), were considerably more abundant in Anamalai, and several major frugivore groups, especially the smaller-bodied bulbuls, bluebirds, barbets, and mynas, were also more abundant there. This pattern is consistent with the fruit resource tracking hypothesis, whereby frugivores aggregate in areas of high fruit availability (Rey, 1995; Burns, 2004). More broadly, the positive association between fruit availability and frugivore abundance is consistent with bottom-up regulation, whereby variation in resource availability can influence consumer abundance (Terborgh, 1986; Morante-Filho et al., 2018). Similar positive relationships between fruit and frugivore abundance have been documented in several tropical assemblages (Blendinger et al., 2012; Mulwa et al., 2013; Naniwadekar et al., 2015), and align with the long-standing view that fruit availability can limit frugivore populations, particularly during periods of resource scarcity (Terborgh, 1986). Thus, resource availability appears to influence frugivore abundance in the direction we anticipated.

Despite supporting higher local abundances of several frugivore groups, Anamalai did not exhibit higher visitation rates on individual fruiting trees. Instead, visitation rates of several frugivore groups were greater in Namdapha. One possible explanation of the observed pattern could be due to resource dilution, whereby high resource availability (up to six times higher in Anamalai compared to Namdapha) allows consumers to distribute their foraging efforts across the larger resource landscape (Otway et al., 2005; Hambäck et al., 2014), reducing visitation rates to any individual fruiting tree despite higher overall frugivore abundances. Conversely, when resources in the local landscape are relatively scarce, as in Namdapha, individual fruiting trees may become relatively more valuable resources, attracting more concentrated visits from avian frugivores. Similar patterns have been reported in which high neighbourhood fruit abundance reduced visitation or fruit removal from focal plants owing to competition between neighbouring fruiting plants and the focal plant (Saracco et al., 2005; Smith & McWilliams, 2014; Hopson et al., 2020; Chenevert et al., 2026). However, these studies were largely confined to fine-scale neighbourhood-level effects or to conspecific fruit densities surrounding focal plants. Our results suggest that an analogous dilution process may operate at substantially broader spatial scales, extending from fruiting neighbourhoods to entire landscapes or communities differing in overall fruit availability.

Visitation rates of individual frugivore species were positively associated with their local abundances in Anamalai but not in Namdapha, suggesting that encounter probability can be an important determinant of interaction frequency (Velho et al., 2012), but that its influence is context-dependent. Our findings for Anamalai are consistent with the expectations under a neutral scenario, whereby common species are likely to contribute to interaction networks disproportionately simply because they encounter potential interaction partners more frequently than rare species (Vázquez et al., 2007). Similar positive relationships between consumer abundance and interaction frequency have been reported in both pollination and seed-dispersal systems (Vizentin-Bugoni et al., 2018; Acevedo-Quintero et al., 2020; Vitorino et al., 2022). Notably, frugivore abundance explained a substantial proportion of variation in visitation rates in Anamalai despite the reversal of the relationship between frugivore abundance and visitation observed across sites. Therefore, while the larger context of resource availability may modulate the overall frugivore visitation on individual fruiting trees, the relative contribution of individual frugivore species to those visitations may or may not be governed by their local abundance. The absence of a similar abundance–visitation relationship in Namdapha, however, indicates that factors beyond local abundance may become more important where fruit resources are relatively scarce, potentially including the distribution and crop size of individual fruiting trees and frugivore movement behaviour (Morales et al., 2013; Naniwadekar et al., 2015; Wolter et al., 2026). Previous studies have also shown frugivore visitation rates to be influenced by fruit crop size and fruit and seed traits (Ramaswami et al., 2019; Naniwadekar et al., 2025; Mandal et al., in press).

Beyond local abundance, we found no evidence that the degree of frugivory influenced visitation rates in either site. Although highly frugivorous species might be expected to visit fruiting trees more frequently because of their greater dependence on fruits, whereas less frugivorous species may visit fruiting trees opportunistically while foraging for alternative resources such as invertebrates (Grass et al., 2014; Pizo et al., 2021), our results did not support this expectation. The absence of a detectable relationship may reflect differences among species in the spatial scales at which they track fruit resources, with sympatric frugivores (even though with a similar degree of frugivory) responding to fruit availability from individual fruiting trees to larger forest patches (Naniwadekar et al., 2015). Moreover, factors not included in our analyses, such as fine-scale resource distribution, frugivore movement behaviour, or seasonal shifts in diet, may play a more important role in shaping visitation patterns, as has been reported elsewhere (Blake & Loiselle, 1991; Côrtes & Uriarte, 2013; Dehling et al., 2022). Thus, while dietary dependence on fruits may influence foraging behaviour, degree of frugivory alone appears insufficient to explain interspecific variation in visitation rates in our study system.

The site-specific differences in the relationship between visitation rates and frugivore abundances have broader implications for studies that use visitation rates as proxies for frugivore abundance. Visitation frequencies are often assumed to reflect local consumer abundance and have therefore been used as indicators of frugivore populations in studies of plant–frugivore interactions (Luck & Daily, 2003; Menke et al., 2012; Albrecht et al., 2013). Our results suggest that this might not always be a valid assumption. Although frugivore visitation rates were positively associated with local abundances in Anamalai, the absence of such a relationship in Namdapha suggests that visitation rates on individual fruiting trees may not be associated with frugivore abundance. Therefore, future studies aiming to estimate frugivore abundance using visitation rates should also consider other factors such as resource context at a larger spatial scale.

Existing phenological information from and around the study regions suggests that peak fruiting coincides with the onset of the monsoon, and that our sampling period coincided substantially with the lean fruiting season at both sites (Datta et al., 2025; Madhavan et al., 2026), when fruit availability may exert particularly strong constraints on frugivore populations and foraging decisions (Chapman et al., 2005). Consistent with this expectation, the site with lower fruit availability, i.e., Namdapha, also supported lower frugivore abundances, in line with predictions of bottom-up regulation. However, because our study represents a single-season comparison, multi-year and year-round monitoring would be necessary (although challenging in wet tropical forests) to determine whether the observed differences in fruit abundance and frugivore populations, and their visitation rates, persist across the full annual cycle, and whether the strength and direction of these relationships change with fluctuations in resource availability.

Despite limitations, our results have important implications for seed dispersal ecology. Most of the avian frugivore groups examined in this study are recognised as key seed dispersers in the Asian wet tropics (Naniwadekar et al., 2019; Gopal et al., 2020). Visitation rate represents one of the main quantitative components of seed dispersal effectiveness (Schupp et al., 2010), and our findings indicate that fruiting trees in landscapes with lower frugivore abundances and lower fruit resources might benefit in terms of the number of visits they receive. Whether the higher visitation rates observed in Namdapha translate into comparable or even greater seed dispersal effectiveness will depend on additional processes such as fruit removal per visit, seed handling behaviour, dispersal distances, and post-dispersal seed fate (Schupp, 1993; Schupp et al., 2010; Nevo et al., 2023). Comparative studies quantifying these downstream components of dispersal effectiveness across resource-rich and resource-poor landscapes would therefore represent a promising avenue for future research.

## ACKNOWLEDGEMENT

We thank the Arunachal Pradesh and Tamil Nadu Forest Departments for granting us the necessary permits to conduct fieldwork in Namdapha and Anamalai Tiger Reserves, respectively. We thank Dhan Bahadur Limbu, Wanggao Wangsa, Laiphong Wangnao, Krishnakumar and Sathiyaraj for assisting in the field in Namdapha and Anamalai, respectively. We thank Vignesh Chandran, Mandar Tijare, Siddhant Mehendale, Prabhav Benara, and Sartaj Ghuman for assisting with data collection at the respective field sites. We thank Divya Mudappa, T. R. Shankar Raman, Navendu Page, Aparajita Datta, Mousumi Ghosh, and M. Ananda Kumar for valuable feedback. We thank Japang Pansa, Srinivasan Kasinathan, Sathish, Stella, and Anamalai Rainforest Research Station for logistical support in the respective field sites.

## REFERENCES

Acevedo-Quintero, J. F., Zamora-Abrego, J. G., & García, D. (2020). From structure to function in mutualistic interaction networks: Topologically important frugivores have greater potential as seed dispersers. Journal of Animal Ecology, 89(9), 2181–2191. 10.1111/1365-2656.13273

Albrecht, J., Berens, D. G., Blüthgen, N., Jaroszewicz, B., Selva, N., & Farwig, N. (2013). Logging and forest edges reduce redundancy in plant–frugivore networks in an old-growth European forest. Journal of Ecology, 101(4), 990–999. 10.1111/1365-2745.12105

Bascompte, J., & Jordano, P. (2007). Plant-animal mutualistic networks: the architecture of biodiversity. *Annual Review of Ecology*, Evolution, and Systematics, 38, 567–593. 10.1146/annurev.ecolsys.38.091206.095818

Bates, D., Mächler, M., Bolker, B., & Walker, S. (2015). Fitting linear mixed-effects models using lme4. Journal of Statistical Software, 67, 1–48. 10.18637/jss.v067.i01

Blake, J. G., & Loiselle, B. A. (1991). Variation in resource abundance affects capture rates of birds in three lowland habitats in Costa Rica. The Auk, 108(1), 114–130. 10.1093/auk/108.1.114

Blendinger, P. G., Loiselle, B. A., & Blake, J. G. (2008). Crop size, plant aggregation, and microhabitat type affect fruit removal by birds from individual melastome plants in the Upper Amazon. Oecologia, 158(2), 273–283. 10.1007/s00442-008-1146-3

Blendinger, P. G., Ruggera, R. A., Montellano, M. G. N., Macchi, L., Zelaya, P. V., Álvarez, M. E., Martín, E., Acosta, O. O., Sánchez, R., & Haedo, J. (2012). Fine-tuning the fruit-tracking hypothesis: Spatiotemporal links between fruit availability and fruit consumption by birds in Andean mountain forests. Journal of Animal Ecology, 81(6), 1298–1310. 10.1111/j.1365-2656.2012.02011.x

Burns, K. C. (2004). Scale and macroecological patterns in seed dispersal mutualisms. Global Ecology and Biogeography, 13(4), 289–293. 10.1111/j.1466-822X.2004.00108.x

Burns, K. C. (2013). What causes size coupling in fruit–frugivore interaction webs? Ecology, 94(2), 295–300. 10.1890/12-1161.1

Chapman, C. A., Chapman, L. J., Zanne, A. E., Poulsen, J. R., & Clark, C. J. (2005). A 12-Year Phenological Record of Fruiting: Implications for Frugivore Populations and Indicators of Climate Change. In J. L. Dew & J. P. Boubli (Eds.), Tropical Fruits and Frugivores: The Search for Strong Interactors (pp. 75–92). Springer Netherlands. 10.1007/1-4020-3833-X_5

Chenevert, C. R., Lamperty, T., Cabrera, D., & Karubian, J. (2026). Effects of fruiting neighborhood, plant traits, and fruit traits on frugivore visitation and fruit removal for an understory palm. Biotropica, 58(2), e70172. 10.1111/btp.70172

Côrtes, M. C., & Uriarte, M. (2013). Integrating frugivory and animal movement: A review of the evidence and implications for scaling seed dispersal. Biological Reviews, 88(2), 255–272. 10.1111/j.1469-185X.2012.00250.x

Datta, A., Banerjee, S., Naniwadekar, R., Thapa, K., Rathore, A., Thapa, K., Brah, T., Nabum, T., Mogar, N., Shukla, U., Sidhu, S., & Borawake, N. (2025). Patterns of leaf, flower, and fruit phenology and environmental relationships in a seasonal tropical forest in the Indian Eastern Himalaya. Biotropica, 57(3), e70030. 10.1111/btp.70030

Davidar, P., & Morton, E. S. (1986). The relationship between fruit crop sizes and fruit removal rates by birds. Ecology, 67(1), 262–265. 10.2307/1938529

Dehling, D. M., Dalla Riva, G. V., Hutchinson, M. C., & Stouffer, D. B. (2022). Niche packing and local coexistence in a megadiverse guild of frugivorous birds are mediated by fruit dependence and shifts in interaction frequencies. The American Naturalist, 199(6), 855–868. 10.1086/718684

Dombroskie, S. L., Tracey, A. J., & Aarssen, L. W. (2016). Leafing intensity and the fruit size/number trade-off in woody angiosperms. Journal of Ecology, 104(6), 1759–1767. 10.1111/1365-2745.12622

Donoso, I., García, D., Martínez, D., Tylianakis, J. M., & Stouffer, D. B. (2017). Complementary effects of species abundances and ecological neighborhood on the occurrence of fruit-frugivore interactions. Frontiers in Ecology and Evolution, 5. 10.3389/fevo.2017.00133

Duffy, J. E. (2002). Biodiversity and ecosystem function: The consumer connection. Oikos, 99(2), 201–219. 10.1034/j.1600-0706.2002.990201.x

Fox, J., Weisberg, S., Price, B., Adler, D., Bates, D., Baud-Bovy, G., Bolker, B., Ellison, S., Firth, D., Friendly, M., Gorjanc, G., Graves, S., Heiberger, R., Krivitsky, P., Laboissiere, R., Maechler, M., Monette, G., Murdoch, D., Nilsson, H., … R-Core. (2026). *car: Companion to Applied Regression* (Version 3.1-5) [Computer software]. https://cran.r-project.org/web/packages/car/index.html

Gopal, A., Mudappa, D., Raman, T. R. S., & Naniwadekar, R. (2020). Forest cover and fruit crop size differentially influence frugivory of select rainforest tree species in Western Ghats, India. Biotropica, 52(5), 871–883. 10.1111/btp.12810

Grass, I., Berens, D. G., & Farwig, N. (2014). Guild-specific shifts in visitation rates of frugivores with habitat loss and plant invasion. Oikos, 123(5), 575–582. 10.1111/j.1600-0706.2013.00920.x

Guerra, T. J., Dayrell, R. L. C., Arruda, A. J., Dáttilo, W., Teixido, A. L., Messeder, J. V. S., & Silveira, F. A. O. (2017). Intraspecific variation in fruit–frugivore interactions: Effects of fruiting neighborhood and consequences for seed dispersal. Oecologia, 185(2), 233–243. 10.1007/s00442-017-3943-z

Hambäck, P. A., Inouye, B. D., Andersson, P., & Underwood, N. (2014). Effects of plant neighborhoods on plant–herbivore interactions: Resource dilution and associational effects. Ecology, 95(5), 1370–1383. 10.1890/13-0793.1

Hopson, R. M., Wilcox, R. C., & Tarwater, C. E. (2020). Frugivores vary in their response to neighborhood effects in a novel ecosystem. Acta Oecologica, 108, 103641. 10.1016/j.actao.2020.103641

Howe, H. F. (1977). Bird activity and seed dispersal of a tropical wet forest tree. Ecology, 58(3), 539–550. 10.2307/1939003

Jordano, P. (2000). Fruits and frugivory. In Seeds: The ecology of regeneration in plant communities (pp. 125–165). 10.1079/9780851994321.0125

Lenth, R. V., Piaskowski, J., Banfai, B., Bolker, B., Buerkner, P., Giné-Vázquez, I., Hervé, M., Jung, M., Love, J., Miguez, F., Riebl, H., Singmann, H., Derumigny, A., & Schmidt, P. (2026). *emmeans: Estimated Marginal Means, aka Least-Squares Means* (Version 2.0.4) [Computer software]. https://cran.r-project.org/web/packages/emmeans/index.html

Luck, G. W., & Daily, G. C. (2003). Tropical countryside bird assemblages: richness, composition, and foraging differ by landscape context. Ecological Applications, 13(1), 235–247. 10.1890/1051-0761(2003)013%5B0235:TCBARC%5D2.0.CO;2

Madhavan, A. P., Kasinathan, S., Murali, A., Sonia, K. B., Moorthi, G., Sundarraj, T., Rajesh, R., Mudappa, D., & Raman, T. R. S. (2026). Phenology of 50 tree species across 9 years in a South Asian tropical rainforest indicates complex influence of climate, traits, and phylogeny (p. 2026.02.14.705701). bioRxiv. 10.64898/2026.02.14.705701

Mandal, R., Gopal, A., Jayanth, A., Sriprasertsil, V., Chaplod, S., Lad, H., Gadkari, A., Desai, N., Kadam, R., Osuri, M.O., Ghuman, S., Page, N., Strange, B.C., Chimchome, V., Joshi, J., & Naniwadekar, R. (in press). Avian frugivore species richness influences dietary breadth and network properties in Asian wet tropical forests. Oikos.

Menke, S., Böhning-Gaese, K., & Schleuning, M. (2012). Plant–frugivore networks are less specialized and more robust at forest–farmland edges than in the interior of a tropical forest. Oikos, 121(10), 1553–1566. 10.1111/j.1600-0706.2011.20210.x

Morales, J. M., García, D., Martínez, D., Rodriguez-Pérez, J., & Herrera, J. M. (2013). Frugivore behavioural details matter for seed dispersal: A multi-species model for Cantabrian thrushes and trees. PLOS ONE, 8(6), e65216. 10.1371/journal.pone.0065216

Morán-López, T., Campagna, M. S., Schleuning, M., García, D., & Morales, J. M. (2025). Global trends of trait matching in avian frugivory and its consequences for the complementarity and irreplaceability of birds. Ecology Letters, 28(2), e70078. 10.1111/ele.70078

Morante-Filho, J., Arroyo-Rodríguez, V., Pessoa, M., Cazetta, E., & Faria, D. (2018). Direct and cascading effects of landscape structure on tropical forest and non-forest frugivorous birds. Ecological Applications, 28. 10.1002/eap.1791

Mulwa, R. K., Neuschulz, E. L., Böhning-Gaese, K., & Schleuning, M. (2013). Seasonal fluctuations of resource abundance and avian feeding guilds across forest–farmland boundaries in tropical Africa. Oikos, 122(4), 524–532. 10.1111/j.1600-0706.2012.20640.x

Muthuramkumar, S., Ayyappan, N., Parthasarathy, N., Mudappa, D., Raman, T. R. S., Selwyn, M. A., & Pragasan, L. A. (2006). Plant community structure in tropical rain forest fragments of the Western Ghats, India. Biotropica, 38(2), 143–160. 10.1111/j.1744-7429.2006.00118.x

Naniwadekar, R., Chaplod, S., Datta, A., Rathore, A., & Sridhar, H. (2019). Large frugivores matter: Insights from network and seed dispersal effectiveness approaches. Journal of Animal Ecology, 88(8), 1250–1262. 10.1111/1365-2656.13005

Naniwadekar, R., Gopal, A., Mandal, R., Jayanth, A., Sriprasertsil, V., Ghuman, S., Page, N., Chaplod, S., Lad, H., Gadkari, A., Chandran, V., Desai, N. A., Kadam, R., Strange, B. C., Chimchome, V., Gale, G. A., & Joshi, J. (2025). Effects of frugivore species pool and seed size on the diversity and functional composition of frugivores visiting fruiting trees. Proceedings of the Royal Society B: Biological Sciences, 292(2056), 20251775. 10.1098/rspb.2025.1775

Naniwadekar, R., Mishra, C., & Datta, A. (2015). Fruit resource tracking by hornbill species at multiple scales in a tropical forest in India. Journal of Tropical Ecology, 31(6), 477–490. 10.1017/S0266467415000449

Naniwadekar, R., Rathore, A., Shukla, U., Chaplod, S., & Datta, A. (2019). How far do Asian forest hornbills disperse seeds? Acta Oecologica, 101, 103482. 10.1016/j.actao.2019.103482

Nevo, O., Filla, C., Valenta, K., & Schupp, E. W. (2023). What drives seed dispersal effectiveness? Ecology and Evolution, 13(9), e10459. 10.1002/ece3.10459

Olson, D. M., Dinerstein, E., Wikramanayake, E. D., Burgess, N. D., Powell, G. V. N., Underwood, E. C., D’amico, J. A., Itoua, I., Strand, H. E., Morrison, J. C., Loucks, C. J., Allnutt, T. F., Ricketts, T. H., Kura, Y., Lamoreux, J. F., Wettengel, W. W., Hedao, P., & Kassem, K. R. (2001). Terrestrial ecoregions of the world: A mew map of life on Earth: A new global map of terrestrial ecoregions provides an innovative tool for conserving biodiversity. BioScience, 51(11), 933–938. 10.1641/0006-3568(2001)051%5B0933:TEOTWA%5D2.0.CO;2

Otway, S. J., Hector, A., & Lawton, J. H. (2005). Resource dilution effects on specialist insect herbivores in a grassland biodiversity experiment. Journal of Animal Ecology, 74(2), 234–240.

Peña, R., Schleuning, M., Dalerum, F., Donoso, I., Rodríguez-Pérez, J., & García, D. (2023). Abundance and trait-matching both shape interaction frequencies between plants and birds in seed-dispersal networks. Basic and Applied Ecology, 66, 11–21. 10.1016/j.baae.2022.11.008

Pizo, M. A., Fontanella, A. B., Carlo, T. A., & González-Castro, A. (2022). Abundance predominates over niche factors as determinant of the frequency of interactions between frugivorous birds and plants. Biotropica, 54(3), 627–634. 10.1111/btp.13076

Pizo, M. A., Morales, J. M., Ovaskainen, O., & Carlo, T. A. (2021). Frugivory specialization in birds and fruit chemistry structure mutualistic networks across the Neotropics. The American Naturalist, 197(2), 236–249. 10.1086/712381

Polis, G. A. (1999). Why are parts of the world green? Multiple factors control productivity and the distribution of biomass. Oikos, 86(1), 3–15. 10.2307/3546565

Proctor, J., Haridasan, K., & Smith. (1998). How far north does Lowland Evergreen Tropical Rain Forest go? Global Ecology & Biogeography Letters, 7(2), 141–146. 10.1111/j.1466-8238.1998.00270.x

Ragusa-Netto, J. (2002). Fruiting phenology and consumption by birds in *Ficus calyptroceras* (Miq.) Miq. (Moraceae). Brazilian Journal of Biology, 62, 339–346. 10.1590/S1519-69842002000200018

Ramaswami, G., Santharam, B., & Quader, S. (2019). Focal plant and neighbourhood fruit crop size effects on fruit removal by frugivores in a semi-arid landscape invaded by *Lantana camara* L. Current Science, 116, 405–411. 10.18520/cs/v116/i3/405-411

Rey, P. J. (1995). Spatio-temporal variation in fruit and frugivorous bird abundance in olive orchards. Ecology, 76(5), 1625–1635. 10.2307/1938163

Saracco, J. F., Collazo, J. A., Groom, M. J., & Carlo, T. A. (2005). Crop size and fruit neighborhood effects on bird visitation to fruiting *Schefflera morototoni* trees in Puerto Rico. Biotropica, 37(1), 81–87. 10.1111/j.1744-7429.2005.04040.x

Schleuning, M., Fründ, J., & García, D. (2015). Predicting ecosystem functions from biodiversity and mutualistic networks: An extension of trait-based concepts to plant–animal interactions. Ecography, 38(4), 380–392. 10.1111/ecog.00983

Schupp, E. W. (1993). Quantity, quality and the effectiveness of seed dispersal by animals. Vegetatio, 107(1), 15–29. 10.1007/BF00052209

Schupp, E. W., Jordano, P., & Gómez, J. M. (2010). Seed dispersal effectiveness revisited: A conceptual review. New Phytologist, 188(2), 333–353. 10.1111/j.1469-8137.2010.03402.x

Sengupta, A. (2022). Animal-mediated seed dispersal in India: Implications for conservation of India’s biodiversity. Biotropica, 54(6), 1320–1330. 10.1111/btp.12982

Smith, A. D., & McWilliams, S. R. (2014). Fruit removal rate depends on neighborhood fruit density, frugivore abundance, and spatial context. Oecologia, 174(3), 931–942. 10.1007/s00442-013-2834-1

Terborgh, J. (1986). Community aspects of frugivory in tropical forests. In A. Estrada & T. H. Fleming (Eds.), Frugivores and seed dispersal (pp. 371–384). Springer Netherlands. 10.1007/978-94-009-4812-9_32

Tilman, D. (1982). Resource Competition and Community Structure. Princeton University Press.

Tobias, J. A., Sheard, C., Pigot, A. L., Devenish, A. J. M., Yang, J., Sayol, F., Neate-Clegg, M. H. C., Alioravainen, N., Weeks, T. L., Barber, R. A., Walkden, P. A., MacGregor, H. E. A., Jones, S. E. I., Vincent, C., Phillips, A. G., Marples, N. M., Montaño-Centellas, F. A., Leandro-Silva, V., Claramunt, S., … Schleuning, M. (2022). AVONET: Morphological, ecological and geographical data for all birds. Ecology Letters, 25(3), 581–597. 10.1111/ele.13898

Vázquez, D. P., Melián, C. J., Williams, N. M., Blüthgen, N., Krasnov, B. R., & Poulin, R. (2007). Species abundance and asymmetric interaction strength in ecological networks. Oikos, 116(7), 1120–1127. 10.1111/j.0030-1299.2007.15828.x

Velho, N., Ratnam, J., Srinivasan, U., & Sankaran, M. (2012). Shifts in community structure of tropical trees and avian frugivores in forests recovering from past logging. Biological Conservation, 153, 32–40. 10.1016/j.biocon.2012.04.028

Vitorino, B. D., Frota, A. V. B. D., Nunes, J. R. D. S., Carniello, M. A., Castrillon, S. K. I., & Maruyama, P. K. (2022). Abundance determines core–periphery roles of birds in a Neotropical seed dispersal network. Ibis, 164(4), 1132–1144. 10.1111/ibi.13075

Vizentin-Bugoni, J., Maruyama, P. K., de Souza, C. S., Ollerton, J., Rech, A. R., & Sazima, M. (2018). Plant-pollinator networks in the tropics: A review. In W. Dáttilo & V. Rico-Gray (Eds.), Ecological Networks in the Tropics: An Integrative Overview of Species Interactions from Some of the Most Species-Rich Habitats on Earth (pp. 73–91). Springer International Publishing. 10.1007/978-3-319-68228-0_6

Wheelwright, N. T. (1985). Fruit size, gape width, and the diets of fruit-eating birds. Ecology, 66(3), 808–818. 10.2307/1940542

White, E. P., Ernest, S. K. M., Kerkhoff, A. J., & Enquist, B. J. (2007). Relationships between body size and abundance in ecology. Trends in Ecology & Evolution, 22(6), 323–330. 10.1016/j.tree.2007.03.007

Wilman, H., Belmaker, J., Simpson, J., Rosa, C. de la, Rivadeneira, M. M., & Jetz, W. (2014). EltonTraits 1.0: Species-level foraging attributes of the world’s birds and mammals. Ecology, 95(7), 2027–2027. 10.1890/13-1917.1

Wolter, J. H., Vissoto, M., Ramírez-Mejía, A. F., Vizentin-Bugoni, J., & Dias, R. A. (2026). Intrinsic and extrinsic drivers shape frugivore functional diversity within a tree population. Ecosphere, 17(2), e70453. 10.1002/ecs2.70453

Whitmore, T. C. (1984). Tropical rain forests of the Par East. Clarendon Press, Oxford. DOI, 10, 0143-6228.

