## Supplementary Material for "More fruits, more frugivores, but not necessarily more visits: findings from two tropical plant–avian frugivore communities"

**Table of contents:**

| **Content** | **Page no.** |
| --- | --- |
| **Table S1.** Frugivore species information | 2–5 |
| **Table S2.** LMM summaries: fruit abundance vs. study site | 5–6 |
| **Table S3.** Pairwise post-hoc comparisons: fruit abundance between study sites | 7 |
| **Table S4:** GLMM summaries: Frugivore encounter rates vs. study sites | 7–8 |
| **Table S5:** Linear model summaries: Frugivore visitation rates vs. study sites | 8–9 |
| **Table S6:** Assessment of multicollinearity among predictor variables included in the LMs | 10 |

| | |
|---|---|
| **Figure S1.** Frugivore encounter rate comparison between morning and evening in Namdapha. | 11 |
| **Figure S2.** Relationship between flock-level and individual-level encounter rates | 12 |

**Table S1.** Avian frugivore species included in the analyses across the two study sites. For each study site, we provide the bird families/frugivore groups, common names and scientific names following Clement's taxonomy for species recorded on transects and/or during focal tree watches.

| Study site | Bird family/frugivore group | Common name | Scientific name (Clement's taxonomy) |
|---|---|---|---|
| Anamalai | Bucerotidae | Great Hornbill | *Buceros bicornis* |
| Anamalai | Bucerotidae | Malabar Grey Hornbill | *Ocyceros griseus* |
| Anamalai | Chloropseidae | Golden-fronted Leafbird | *Chloropsis aurifrons* |
| Anamalai | Columbidae | Grey-fronted Green-Pigeon | *Treron affinis* |
| Anamalai | Columbidae | Malabar Imperial-Pigeon | *Ducula cuprea* |
| Anamalai | Corvidae | White-bellied Treepie | *Dendrocitta leucogastra* |

| | | | |
|---|---|---|---|
| Anamalai | Dicaeidae | Nilgiri Flowerpecker | *Dicaeum concolor* |
| Anamalai | Dicaeidae | Thick-billed Flowerpecker | *Pachyglossa agilis* |
| Anamalai | Irenidae | Asian Fairy-bluebird | *Irena puella* |
| Anamalai | Leiothrichidae | Brown-cheeked Fulvetta | *Alcippe poioicephala* |
| Anamalai | Megalaimidae | Malabar Barbet | *Psilopogon malabaricus* |
| Anamalai | Megalaimidae | White-cheeked Barbet | *Psilopogon viridis* |
| Anamalai | Muscicapidae | Malabar Whistling-Thrush | *Myophonus horsfieldii* |
| Anamalai | Oriolidae | Indian Golden Oriole | *Oriolus kundoo* |
| Anamalai | Psittaculidae | Malabar Parakeet | *Psittacula columboides* |
| Anamalai | Psittaculidae | Plum-headed Parakeet | *Psittacula cyanocephala* |
| Anamalai | Psittaculidae | Vernal Hanging-Parrot | *Loriculus vernalis* |
| Anamalai | Pycnonotidae | Flame-throated Bulbul | *Rubigula gularis* |
| Anamalai | Pycnonotidae | Yellow-browed Bulbul | *Acritillas indica* |
| Anamalai | Pycnonotidae | Gray-headed Bulbul | *Microtarsus priocephalus* |
| Anamalai | Pycnonotidae | Red-whiskered Bulbul | *Pycnonotus jocosus* |
| Anamalai | Sturnidae | Southern Hill Myna | *Gracula indica* |
| Anamalai | Sturnidae | Malabar Starling | *Sturnia blythii* |
| Anamalai | Trogonidae | Malabar Trogon | *Harpactes fasciatus* |
| Anamalai | Turdidae | Orange-headed Thrush | *Geokichla citrina* |
| Namdapha | Bucerotidae | Great Hornbill | *Buceros bicornis* |

| | | | |
|---|---|---|---|
| Namdapha | Bucerotidae | Rufous-necked Hornbill | *Aceros nipalensis* |
| Namdapha | Bucerotidae | Wreathed Hornbill | *Rhyticeros undulatus* |
| Namdapha | Bucerotidae | Brown Hornbill | *Anorrhinus austeni* |
| Namdapha | Campephagidae | Black-winged Cuckooshrike | *Lalage melaschistos* |
| Namdapha | Campephagidae | Oriental Cuckooshrike | *Coracina javensis* |
| Namdapha | Chloropseidae | Blue-winged Leafbird | *Chloropsis moluccensis* |
| Namdapha | Chloropseidae | Orange-bellied Leafbird | *Chloropsis hardwickii* |
| Namdapha | Columbidae | Ashy-headed Green-Pigeon | *Treron phayrei* |
| Namdapha | Columbidae | Mountain Imperial-Pigeon | *Ducula badia* |
| Namdapha | Columbidae | Pin-tailed Green-Pigeon | *Treron apicauda* |
| Namdapha | Irenidae | Asian Fairy-bluebird | *Irena puella* |
| Namdapha | Leiothrichidae | Blue-winged Minla | *Actinodura cyanouroptera* |
| Namdapha | Leiothrichidae | Greater Necklaced Laughingthrush | *Pterorhinus pectoralis* |
| Namdapha | Leiothrichidae | Long-tailed Sibia | *Heterophasia picaoides* |
| Namdapha | Leiothrichidae | Silver-eared Mesia | *Leiothrix argentauris* |
| Namdapha | Leiothrichidae | Beautiful Sibia | *Heterophasia pulchella* |
| Namdapha | Leiothrichidae | Rusty-fronted Barwing | *Actinodura egertoni* |
| Namdapha | Leiothrichidae | Red-tailed Minla | *Minla ignotincta* |
| Namdapha | Leiothrichidae | White-crested | *Garrulax leucolophus* |

| | | | |
|---|---|---|---|
| | | Laughingthrush | |
| Namdapha | Megalaimidae | Blue-throated Barbet | *Psilopogon asiaticus* |
| Namdapha | Megalaimidae | Great Barbet | *Psilopogon virens* |
| Namdapha | Megalaimidae | Blue-eared Barbet | *Psilopogon cyanotis* |
| Namdapha | Megalaimidae | Golden-throated Barbet | *Psilopogon franklinii* |
| Namdapha | Muscicapidae | Blue Whistling-Thrush | *Myophonus caeruleus* |
| Namdapha | Oriolidae | Maroon Oriole | *Oriolus traillii* |
| Namdapha | Picidae | Greater Yellownape | *Chrysophlegma flavinucha* |
| Namdapha | Pycnonotidae | Ashy Bulbul | *Hemixos flavala* |
| Namdapha | Pycnonotidae | White-throated Bulbul | *Alophoixus flaveolus* |
| Namdapha | Pycnonotidae | Black Bulbul | *Hypsipetes leucocephalus* |
| Namdapha | Sturnidae | Common Hill Myna | *Gracula religiosa* |
| Namdapha | Trogonidae | Red-headed Trogon | *Harpactes erythrocephalus* |
| Namdapha | Turdidae | Green Cochoa | *Cochoa viridis* |
| Namdapha | Zosteropidae | Whiskered Yuhina | *Yuhina flavicollis* |
| Namdapha | Zosteropidae | Black-chinned Yuhina | *Yuhina nigrimenta* |

**Table S2.** Results of three separate linear mixed-effects models examining whether log-transformed fruit abundance for small-, medium-, and large-seeded bird-consumed fruiting species was different between our study sites – Anamalai and Namdapha. In each model, we included transect-patch identity as a random intercept.

| Seed size | Fixed effect | Estimate | SE | *t*-value |
|---|---|---|---|---|
| Small | Intercept | 3.91 | 0.47 | 8.29 |
| Small | Site: Namdapha | -1.74 | 0.69 | -2.53 |
| Medium | Intercept | 1.92 | 0.44 | 4.34 |
| Medium | Site: Namdapha | -0.95 | 0.62 | -1.53 |
| Large | Intercept | 1.26 | 0.23 | 5.43 |
| Large | Site: Namdapha | -0.28 | 0.34 | -0.83 |
| Seed size | Random effect | Variance | | SD |
| Small | Patch ID | 1.40 | | 1.18 |
| Small | Residual | 15.01 | | 3.87 |

| | | | |
|---|---|---|---|
| Medium | Patch ID | 2.09 | 1.44 |
| Medium | Residual | 4.44 | 2.11 |
| Large | Patch ID | 0.27 | 0.52 |
| Large | Residual | 4.29 | 2.07 |

**Table S3.** Pairwise post-hoc comparisons of log-transformed fruit abundance between sites within each seed-size category.

| Seed size | Contrast (Anamalai - Namdapha) | *p*-value |
|---|---|---|
| Small | 1.74 | 0.018 |
| Medium | 0.95 | 0.140 |
| Large | 0.28 | 0.415 |

**Table S4:** Results of negative binomial generalised linear mixed-effects models examining differences in frugivore encounter rates between Anamalai and Namdapha for each frugivore group. Estimates and standard errors (SE) are presented on the log scale. Anamalai was the reference site; therefore, the intercept represents the estimated log encounter rate in Anamalai, and the coefficient for Site: Namdapha represents the difference in log encounter rate between Namdapha and Anamalai.

| Frugivore group | Fixed effect | Estimate (log scale) | SE | *z*-score | *p*-value |
|---|---|---|---|---|---|
| Megalaimidae | Intercept | 2.03 | 0.08 | 26.77 | <0.001 |
| Megalaimidae | Site: Namdapha | -0.96 | 0.13 | -7.56 | <0.001 |
| Irenidae | Intercept | 0.93 | 0.10 | 9.71 | <0.001 |
| Irenidae | Site: Namdapha | -1.20 | 0.19 | -6.19 | <0.001 |
| Pycnonotidae | Intercept | 1.64 | 0.06 | 27.29 | <0.001 |
| Pycnonotidae | Site: Namdapha | -0.84 | 0.12 | -7.33 | <0.001 |
| Bucerotidae | Intercept | 0.64 | 0.10 | 6.37 | <0.001 |
| Bucerotidae | Site: Namdapha | 0.06 | 0.15 | 0.37 | 0.713 |
| Sturnidae | Intercept | 0.81 | 0.14 | 5.63 | <0.001 |
| Sturnidae | Site: Namdapha | -2.01 | 0.31 | -6.50 | <0.001 |

| | | | | | |
|---|---|---|---|---|---|
| Columbidae | Intercept | -0.92 | 0.22 | -4.17 | <0.001 |
| Columbidae | Site: Namdapha | 0.28 | 0.32 | 0.89 | 0.372 |

**Table S5:** Summary of the linear models examining the effects of site on log-transformed frugivore visitation rates on fruiting trees for each frugivore group.

| Frugivore group | Fixed effect | Estimate | SE | *p*-value |
|---|---|---|---|---|
| Megalaimidae | Intercept | 0.05 | 0.28 | 0.855 |
| Megalaimidae | Site: Namdapha | -0.37 | 0.41 | 0.370 |
| Irenidae | Intercept | -0.39 | 0.32 | 0.234 |
| Irenidae | Site: Namdapha | 1.28 | 0.62 | 0.048 |
| Pycnonotidae | Intercept | 0.36 | 0.20 | 0.082 |
| Pycnonotidae | Site: Namdapha | 1.04 | 0.32 | 0.003 |
| Bucerotidae | Intercept | -1.27 | 0.35 | 0.001 |
| Bucerotidae | Site: Namdapha | 1.41 | 0.50 | 0.009 |

| | | | | |
|---|---|---|---|---|
| Sturnidae | Intercept | -0.50 | 0.34 | 0.155 |
| Sturnidae | Site: Namdapha | 0.60 | 0.70 | 0.397 |
| Columbidae | Intercept | -0.75 | 0.41 | 0.087 |
| Columbidae | Site: Namdapha | 0.63 | 0.54 | 0.260 |

**Table S6:** Assessment of multicollinearity among predictor variables included in the linear models relating frugivore visitation rates to local abundance (encounter rate), degree of frugivory, and body mass.

| Site | Predictor (scaled) | Variance Inflation Factor (VIF) |
|---|---|---|
| Anamalai | Frugivore encounter rate | 1.01 |
| Anamalai | Degree of frugivory | 1.03 |
| Anamalai | Body mass | 1.04 |
| Namdapha | Frugivore encounter rate | 1.05 |
| Namdapha | Degree of frugivory | 1.27 |

| | | |
|---|---|---|
| Namdapha | Body mass | 1.24 |

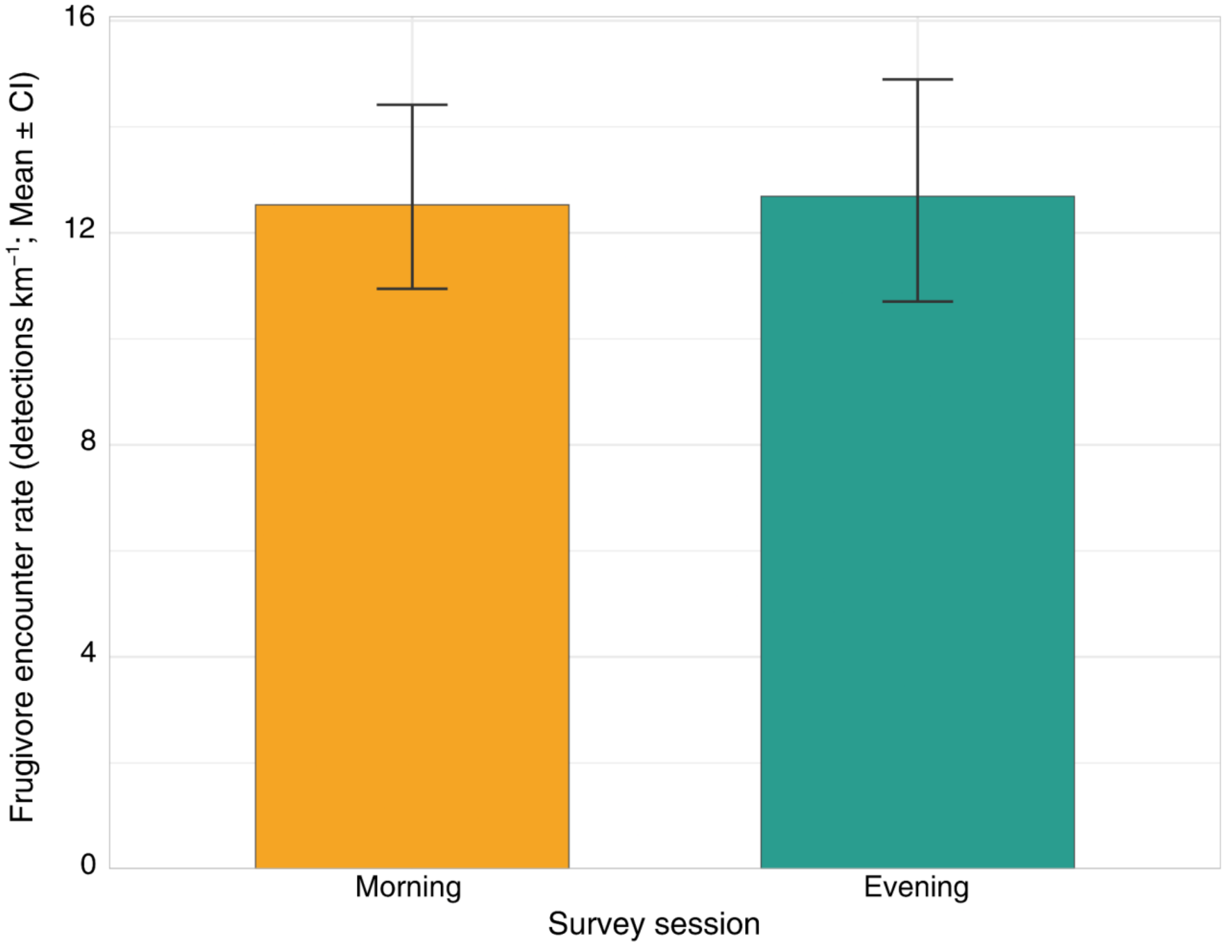

**Figure S1.** Mean encounter rates of avian frugivores during morning and evening survey sessions in Namdapha. Bars represent mean encounter rates (detections km$^{-1}$), and error bars indicate bootstrapped 95% confidence intervals (with 1,000 iterations).

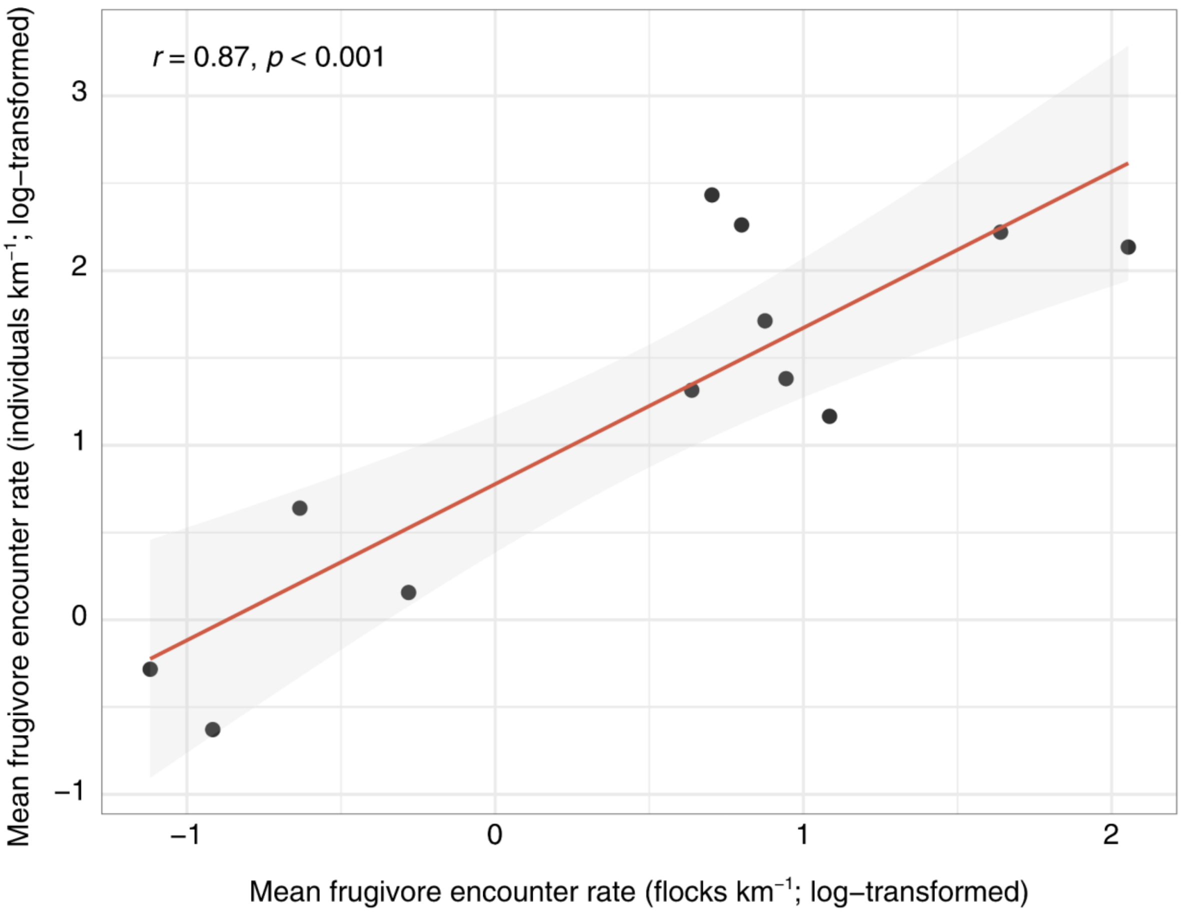

**Figure S2.** Relationship between flock-level and individual-level encounter rates of avian frugivores across the two study sites. Points represent the mean encounter rates for each of the six major frugivore groups in Anamalai and Namdapha (Pycnonotidae, Irenidae, Megalaimidae, Sturnidae, Columbidae, and Bucerotidae). Both encounter rates were log-transformed prior to analysis. The solid line represents the fitted linear regression, and the shaded band indicates the 95% confidence interval. Pearson's correlation coefficient ($r = 0.87$, $p < 0.001$) indicates a strong positive association between flock-level and individual-level encounter rates.
